# Quantitating SARS-CoV-2 Neutralizing Antibodies from Human Dried Blood Spots

**DOI:** 10.1101/2024.03.18.585599

**Authors:** Katherine Berman, Greta Van Slyke, Hayley Novak, Jean M. Rock, Rachel Bievenue, Amanda K. Damjanovic, Kate L. DeRosa, Gianna Mirabile, Roxie C. Girardin, Alan P. Dupuis, Kathleen A. McDonough, Monica M. Parker, Linda M. Styer, Nicholas J. Mantis

## Abstract

**Background:** In the earliest days of COVID-19 pandemic, the collection of dried blood spots (DBS) enabled public health laboratories to undertake population-scale seroprevalence studies to estimate rates of SARS-CoV-2 exposure. With SARS-CoV-2 seropositivity levels now estimated to exceed 94% in the United States, attention has turned to using DBS to assess functional (neutralizing) antibodies within cohorts of interest.

**Methods:** Contrived DBS eluates from convalescent, fully vaccinated and pre-COVID-19 serum samples were evaluated in SARS-CoV-2 plaque reduction neutralization titer (PRNT) assays, a SARS-CoV-2 specific 8-plex microsphere immunoassay, a cell-based pseudovirus assay, and two different spike-ACE2 inhibition assays, an in-house Luminex-based RBD-ACE2 inhibition assay and a commercial real-time PCR-based inhibition assay (NAB-Sure™).

**Results:** DBS eluates from convalescent individuals were compatible with the spike-ACE2 inhibition assays, but not cell-based pseudovirus assays or PRNT. However, the insensitivity of cell-based pseudovirus assays was overcome with DBS eluates from vaccinated individuals with high SARS-CoV-2 antibody titers.

**Conclusion:** SARS-CoV-2 neutralizing titers can be derived with confidence from DBS eluates, thereby opening the door to the use of these biospecimens for the analysis of vulnerable populations and normally hard to reach communities.

## 1. INTRODUCTION

The collection and serologic analysis of dried blood spots (DBS) from population-based serosurveys proved integral to understanding the dynamics of SARS-CoV-2 infection in the first months of the COVID-19 pandemic. In New York State, for example, a statewide seroprevalence study among ∼15,000 individuals in public settings (e.g., grocery store fronts) conducted in April 2020 afforded some of the first insights into overall disease incidence and demographics of virus exposure in urban and rural communities (1). Similarly, a cross-sectional serosurvey of ∼400,000 newborns in New York State that employed heel stick DBS collected between November 2019 and November 2021 afforded unprecedent insight into COVID-19 exposures and vaccination status among a diverse population of reproductive-aged females (2). Other groups have leveraged clinician-assisted and self-collected DBS to conduct large scale serosurveys of vulnerable cohorts, school age children, and otherwise hard to reach communities (3–10). In addition to ease of collection, once collected onto Whatman filter paper, DBS samples are stable for long periods of time under refrigeration (4-8°C) or frozen (up to - 20 °C) in the presence of desiccants (11, 12).

Within the COVID-19 pandemic, DBS have largely been employed in serological studies aimed at determining SARS-CoV-2 S and N reactivity as measures of infection and vaccination. However, with SARS-CoV-2 seropositivity levels now estimated to exceed 94% in the United States, the value of such serosurveys has diminished (13). Furthermore, the emergence of omicron subvariants with highly mutated spike proteins confounds one’s ability to extrapolate neutralizing antibody titers from binding antibody units (BAUs) (14, 15). For those and other reasons, there is considerable interest in developing a method to assess SARS-CoV-2 neutralizing titers directly from DBS eluates.

Indeed, several reports have already described the use of DBS eluates to assess functional activity of SARS-COV-2 antibodies. For example, Itell and colleagues examined paired plasma and fingerstick (FS) DBS from convalescent individuals in SARS-COV-2 spike-pseudotyped lentiviral particle assay in 96- and 384-well format (16). In another report, Sancilio and colleagues successfully used an ACE2-RBD binding assay on the Mesoscale Diagnostics (MSD) platform to estimate neutralizing titers in contrived DBS (cDBS) eluates (17) and in self-collected DBS from a community-based serology survey (18). However, neither study benchmarked their results against plaque reduction neutralization titer (PRNT) values, nor did they perform cross comparisons of the different *in vitro* assays (e.g., pseudovirus versus ACE2-RBD).

Therefore, to further advance the utility and validity of DBS for the purpose of functional SARS-COV-2 serology, we investigated the compatibility of contrived DBS eluates with different SARS-COV-2 neutralization assays and compared their outputs to PRNT values derived from matched serum. While DBS eluates proved incompatible with cell-based SARS-COV-2 pseudovirus neutralization assays, we found that two different ACE2 inhibition assays, an in-house Luminex-based assay, and a commercial real-time PCR-based assay (NAB-Sure™), yielded neutralization values that strongly correlated with PRNT. An 8-plex MIA based solely on binding antibodies to spike antigens also served as a proxy for PRNT. As the two RBD-ACE2 inhibition assays employed in this study are amenable to high-throughput testing, we propose to integrate one or both into our DBS workflow for analysis of cohort specific studies going forward.

## 2. METHODS

### 2.1 SARS-CoV-2 reagents

The source and descriptions of SARS-CoV-2 reagents used in this study is provided in **Table 1**.

**Table 1.** SARS-CoV-3 Reagents and Sources.

| **Table 1. SARS-CoV-3 Reagents and Sources** |  |  |  |  |
| --- | --- | --- | --- | --- |
| **Reagent** | **Description** | **Source** | **Catalogue Number** | **Usage** |
| TBS | Tris Buffered Saline with Casein | BioRad | 1610782 | DBS Elution |
| FLT | Full Length Trimeric SARS-CoV-2 Spike | MassBiologics | N/A | 8-Plex MIA/iACE2 |
| FLS | Full length SARS-CoV-2 Spike, His-Tag | Native Antigen | REC31868-100 | 8-Plex MIA |
| Spike Subunit 1 (S1) | SARS-CoV-2 (2019-nCoV) Spike S1, His-Tag | Sino Biological | 40591-V08H | 8-Plex MIA |
| RBD | SARS-CoV-2 Stable RBD, Thrombin-His | MassBiologics | N/A | 8-Plex MIA |
| Nucleocapsid (N) | 2019-nCoV Nucleocapsid Protein, His tag | SinoBiological | 40588-V08B | 8-Plex MIA |
| Nucleocapsid (N-NA) | SARS-CoV-2 Nucleoprotein, His-Tag (E. coli) | Native Antigen | REC31812-100 | 8-Plex MIA |
| Nucleocapsid (NHT) | SARS-CoV-2 (2019-nCoV) Nucleocapsid Protein (His tag) | Sino Biological | 0588-V07E | 8-Plex MIA |
| Internal Control (IC) | Mouse anti-Human IgG3 Secondary Antibody | ThermoFisher | MA1-83242 | 8-Plex MIA |
| WT sRBD | SARS (WA1/2020) RBD, Thrombin-His | MassBiologics | N/A | iACE2 |
| hACE2 | Biotinylated ACE2 | Wadsworth Center Protein Core | N/A | iACE2 |
| CC12.3 | Human Mab RBD-A WT D614G | Scripps | N/A | iACE2/RVP |
| RVP-702L | WT D614G Strain Reporter Virus Particles | Integral Molecular | CL-275A | RVP |
| 293-hsACE2 Cells | 293T cells expressing human ACE2 receptor | Integral Molecular | TA-060520-MC | RVP |

### 2.2 Human serum samples

A set of serum samples from 30 PCR positive donors was purchased from Access Biologicals, LLC, (Vista, CA), herein referred to as “Panel D”. Positive test results were reported in the initial U.S. wave of infections between 3/15/2020 and 4/25/2020; and samples were drawn between 6/3/2020-6/4/2020, indicating that all donors were infected with the Washington variant of SARS-CoV-2. The panel consists of 21 female and 9 male donors; ages ranged from 25-to 66-years-old; all donors were symptomatic. A set of 87 COVID-19 negative serum samples collected between 3/24/2017 and 11/9/2018 was also purchased from Access Biologicals, LLC. This set is referred to as “Panel E”, which consists of pre-pandemic SARS-CoV-2 sero-negative donors (73 females and 14 males between 18 and 64-years-old). A third set of samples from 30 COVID-19 vaccinated patients was purchased from Access Biologicals. 15 of these samples were selected for evaluation in this study. Participants were vaccinated with the Moderna COVID-19 vaccine between December 2020 and March 2021. Samples were collected 12-17 days after their second dose. This set is referred to as ‘Panel H’. Upon receipt, Panels D, E and H serum samples were stored at −80°C.

### 2.3 Contrived DBS

Contrived DBS were created by mixing an equal volume of Panel D, E or H serum with washed human blood cells (BC). Fifty microliters (50 μL) of the BC-serum mixture were spotted onto Whatman 903 filter paper (SigmaAldrich, St. Louis, MO) and allowed to air dry at room temperature for 4 h before being packaged in semi-permeable bags with desiccant and stored at −20°C. Blood cells (BC) were collected by centrifugation (2200 RCF for 8 min) from commercially obtained fresh human whole blood [Type O EDTA] (ZenBio, Durham, NC).

### 2.4 Plaque reduction neutralization titer (PRNT)

PRNT was determined as described (19, 20). Briefly, 2-fold serially diluted serum (100 μl) was mixed with an equal volume of 150-200 plaque forming units (PFUs) of SARS-CoV-2 (isolate USA-WA1/2020, BEI Resources, NR-52281) and incubated for 1 h at 37°C, 5% CO_2_. The virus:serum mixture (100 μl) was applied to VeroE6 cells grown to 95%–100% confluency in 6 well plates. Adsorption of the virus:serum mixture was allowed to proceed for 1 h at 37°C, 5% CO_2._ Following the adsorption period, the wells were overlaid with 0.6% agar prepared in cell culture medium (Eagle’s Minimal Essential Medium, 2% heat inactivated FBS, 100 μg/ml penicillin G, 100 U/ml streptomycin). Two days post-infection, a second agar overlay containing 0.2% neutral red prepared in cell culture medium was applied. Plaque determinations were conducted 1-2 days later. Neutralizing titers were defined as the inverse of the highest dilution of serum providing 50% (PRNT50) or 90% (PRNT90) viral plaque reduction relative to a virus only control.

### 2.5 Microsphere Immunoassay (MIA)

The development and validation of an 8-plex MIA consisting of three different nucleocapsid N antigens, RBD, Spike S1 (S1), full length Spike (FLS), and full length trimer (FLT) is described elsewhere (J. Rock, L.Styer, *manuscript in preparation*). Magplex-C microspheres (Luminex Corp) were coupled with 7 different SARS-CoV-2 antigens and 1 internal control antibody. Each antigen/antibody was coupled to a different bead region allowing the beads to be multiplexed in the same well. A 3.3 mm DBS punch from each sample was combined with 250 µl elution buffer (Tris-buffered saline, 1% casein blocker) (Bio-Rad Laboratories) in a 96 well plate. The DBS punches were eluted for 1 hour at room temperature. Antigen-coated beads (diluted to 1,250 beads/bead set/well with assay buffer (PBS, 2% BSA, pH 7.4)) and 25 µl DBS eluate were combined in a 384 well plate and incubated at 37°C for 30 minutes while shaking (300 RPM) in the dark. Plates were washed for two cycles using the BioTek 405TS microplate washer and 50 µl wash buffer (PBS, 2% BSA, 0.02% Tween-20, 0.05% sodium azide, pH 7.4). 50 µl phycoerythrin-tagged goat anti-human IgG was added to each well. Plates were incubated at 37°C for 30 minutes while shaking (300 RPM) in the dark and washed as described above. Antibody-bead complexes were resuspended with 90 µl xMap sheath fluid (Luminex), plates were incubated for 1 minute at room temperature while shaking (300 RPM), and 70 µl from each well was read on a Luminex FlexMAP 3D instrument. Results were presented in median fluorescence intensity (MFI) for each well. Positive, negative, and background controls were included with every plate.

### 2.6 Reporter virus particle (RVP) neutralization assays

Pseudo-typed lentiviral reporter virus particles (RVP) tagged with Renilla luciferase (Luc) and 293T-hsACE2 cells were obtained from Integral Molecular, Inc. (Philadelphia, PA). To determine the SARS-CoV-2 neutralizing activity in human serum and paired cDBS eluates, serum and DBS eluates were diluted 1:50 in DMEM containing 10mM HEPES [pH 7.4], 10% fetal bovine serum and 100 U/mL penicillin and 0.1 mg/mL streptomycin (Sigma-Aldrich). Paired sample types were serially diluted in duplicate in cell medium. RVPs were thawed at 37°C and immediately diluted per the manufacturer’s specification in cell medium and then 50 µl was added to the assay plate. A total of 50 µl of diluted human serum or DBS eluate were then transferred from the dilution plate and mixed with RVP solution. Serum and DBS eluates were incubated with RVPs for 1 hour at 37°C with 5% CO_2_. Positive and negative control wells were included on each assay plate and consisted of RVPs only (100% infectivity control), and no RVPs (0% infectivity control). During RVP-serum incubation 80-90% confluent 293T-hsACE2 cells were harvested and cell density was determined using a hemocytometer. Cells were diluted to a density of 2 x 10^5^ cells/mL in cell culture media and 100 µl was added to each well of the RVP + specimen plate and mixed with gentle pipetting. Plates were incubated at 37°C with 5% CO_2_ for 72 h. Quantification of SARS-CoV-2 RVP infection was determined using the *Renilla*-Glo^®^ luciferase assay system (Promega Corp., Madison, WI). Microtiter assay plates were centrifuged for 5 min at 2000 rpm to prevent cell loss, then supernatant was removed with a pipet and 30 µl of PBS was added to each well. *Renilla*-Glo^®^ substrate diluted 1:100 (v/v) into buffer was prepared fresh; 30 µl added per well and plates were then incubated for 10 min at room temperature. Relative light units (RLU) were detected on a SpectroMax iD3 (Molecular Devices, LLC, San Jose, CA) and analyzed with SoftMax Pro 7.0 software. A Z’ value was calculated for each plate for assay accuracy and comparability, plates with a Z’<0.5 were repeated. The RLU mean and standard deviation (SD) of 100% infected wells was determined, and a minimum limit of detection was calculated for each plate as = (Mean– (3xSD)) _100%Infected_. Samples that failed to reach the LOD cut-off value at the most concentrated dilution were classified as non-neutralizing (<LOD) and assigned a value of 0 for statistical analysis purposes. For each plate the RLU mean value for 0% infected wells was subtracted from the mean of 100% infected wells resulting in a normalized infection value (NIV); % normalized RVP neutralization was calculated as = 100-((S_AVE_/NIV)*100), where S_AVE_= the average of duplicate sample RLU values.

### 2.7 Human ACE2 inhibition (iACE2) assay

Recombinant soluble human ACE2 was generated by the Wadsworth Center’s Protein Expression core facility. The coding region for human ACE2 (residues 1-615; Genbank #) was cloned into pcDNA3-sACE2(WT)-8his [(https://www.addgene.org/browse/sequence/298420/] and transfected into ExpiCHO-S cells using ExpiFectamine, as per manufacturer’s instructions (ThermoFisher (https://www.thermofisher.com/order/catalog/product/A29133). The recombinant protein was affinity purified on 2 x 1ml HisTrap™ Excel (Cytiva) columns connected in series. ACE2 was eluted with a linear gradient of 20 mM Tris-HCl, 500 mM NaCl, 500 mM imidazole pH 8.0 over 20 column volumes. Fractions containing ACE2, as assessed by SDS-PAGE, were pooled and dialyzed twice against PBS [pH 7.4]. Protein concentration assessed by Bradford assay with bovine albumin as a standard curve. Biotinylated using EZ-Link NHS Biotin (Thermo Fisher Scientific).

Magplex-C microspheres (Luminex) from separate bead regions were coupled with RBD and FLT from SARS-CoV-2 isolate USA-WA1/2020 (“wild type”) and resuspended to 2500 beads/region/well. Diluted beads were transferred to wells of a non-binding 96 well plate; 50 µl of serum or DBS eluate (1:138) was then added to each well and incubated for 1 h RT with agitation in the dark. Plates were washed twice manually with wash buffer (190 µl) and then biotinylated-hACE2 (see below; 4 µg/mL) was added to each well, plates were incubated for an additional 45 min at RT, shaking in the dark. Plates were washed, probed with streptavidin-PE (1 µg/mL) and incubated at RT for 30 min. The plates were washed to removed unbound streptavidin-PE then analyzed using FlexMap3D. The MFI for ACE2 binding (in the absence of antibody) was derived from a single column of wells containing diluted beads. Inhibition of ACE2 (iACE2) binding calculated as: 190 µl % ACE2 binding-inhibition (iACE2), per sample well, where % iACE2 = 100-[Raw value/(Average of wells with specific VoC beads and no serum or DBS eluate) x 100].

### 2.8 NAB-Sure™

The NAB-Sure™ SARS-CoV-2 Neutralizing Antibody Test (Spear Bio, MA, USA) was performed according to the manufacturer’s instructions (www.spear.bio/nabsure). NAB-Sure™ is a competitive binding assay that measures the ability of antibodies in DBS samples to block SARS-CoV-2 Spike Protein (S1) from binding to recombinant hACE2 in solution. The assay was validated at the Wadsworth Center using eluates (n=11) from SARS-CoV-2 infected individuals with range of Spike-specific titers provided by SpearBio. Contrived Panel D and H DBS punches (6mm) were incubated in NAB-Sure™ elution buffer at 37°C for 1 h. Afterwards, eluates (15μL) were processed according to the manufacturer’s instructions. The test was read using a 7500 Fast Real-Time PCR System (Thermo Fisher Scientific) in the Wadsworth Center’s Bloodborne Virus Laboratory. CT values were converted to NT_50_ values relative to the negative control sample that is devoid of SARS-CoV-2 neutralizing antibodies. The cycle difference between the sample CT value and the negative control CT value was calculated. Inhibition % was calculated as = 1-1/2^x^, where x= cycle difference. An interpolated NT50 value was determined using GraphPad Prism software. The final NT50 value= y*50, where y=interpolated NT50 value.

### 2.9 Statistical Analysis

Statistical analyses were performed using GraphPad Prism 9 software. Correlations were assessed by the Pearson correlation coefficient. An unpaired, two-tailed Welch’s t-test was performed to determine significance of differences in ACE2 inhibition between the two groups. Statistical significance of differences in RVP dilutions was assessed using a 2-way ANOVA. In all cases, the following apply: *p ≤ 0.05, **p≤ 0.01, ***p≤ 0.001, ****p < 0.0001.

## 3. RESULTS

### 3.1 Generation of standard SARS-CoV-2 DBS panel

The primary objective of this study was to assess SARS-CoV-2 virus neutralizing activity from DBS eluates benchmarked against PRNT assays, since performing actual PRNT assays on DBS eluates is neither practical nor technically feasible. For this purpose, we generated contrived DBS (cDBS) from a commercial panel (“Panel D”) consisting of 30 individuals with a defined history of SARS-CoV-2 infection during or prior to June 2020. The participants ranged in age from 25 to 63 (average 45) with 21 females and 9 males. Of the 30 COVID-19 positive samples, 26 were confirmed positive via SARS-CoV-2 PCR and four strictly through medical diagnosis. PRNT_50_ values for 15 of the 30 samples were determined using SARS-CoV-2 Washington (WA1), as described (19). The average PRNT_50_ for the cohort was 351 (+/- 271). In addition, the percent SARS-CoV-2 neutralization at a fixed 1:100 serum dilution (average 55 +/- 22%) was extrapolated based on PRNT curves to enable direct comparison with values derived from DBS eluates (see below). A summary of the 15 Panel D participants is provided in **Table 2**.

**Table 2.** Panel D Summary.

| **Table 2. Panel D Summary** |  |  |  |  |  |  |  |  |  |  |  |  |  |  |
| --- | --- | --- | --- | --- | --- | --- | --- | --- | --- | --- | --- | --- | --- | --- |
|  |  |  | **Serum (PRNT)** |  | **MFI in 8-plex MIA by Antigen** |  |  |  |  |  | **iACE2** | **RVP** |  | **NAb** |
| **#** | **Age** | **Gender** | **NT<sub>50</sub>** | **%Neut** | **N<sup>a</sup>** | **S1<sup>b</sup>** | **FLS<sup>c</sup>** | **FLT<sup>d</sup>** | **RBD<sup>e</sup>** | **Sum<sup>f</sup>** | **iACE2<sup>g</sup>** | **NT50** | **%Neut** | **NT<sub>50</sub>** |
| 1 | 38 | F | >832 | 93 | 6121.0 | 4915 | 10178 | 14238 | 11185 | 40516 | 43 +/-5.3 | 379 | 89 | 712 |
| 2 | 45 | F | 104 | 51 | 3283.5 | 521 | 1514 | 2557 | 1509 | 6100 | 17 +/-4.3 | 183 | 65 | 90 |
| 3 | 44 | F | 208 | 42 | 2925.5 | 2377 | 4880 | 6843 | 5306 | 19406 | 22 +/-2.2 | 151 | 58 | 244 |
| 4 | 48 | F | >832 | 93 | 3396.0 | 4295 | 9836 | 15808 | 11335 | 41274 | 53 +/-2.7 | 203 | 79 | 557 |
| 5 | 55 | F | 416 | 33 | 544.0 | 872 | 2779 | 4562 | 2868 | 11081 | 14 +/-3.4 | 435 | 67 | 196 |
| 6 | 65 | F | 208 | 60 | 4560.0 | 3221 | 6926 | 7929 | 5972 | 24047 | 14 +/-3.4 | 395 | 64 | 270 |
| 7 | 40 | F | 208 | 77 | 7424.0 | 1714 | 4967 | 6497 | 5496 | 18674 | 18 +/-2.3 | 513 | 67 | 189 |
| 8 | 39 | M | 832 | 70 | 6465.0 | 4561 | 8677 | 16991 | 10578 | 40806 | 42 +/-7.1 | 262 | 71 | 899 |
| 9 | 34 | F | IND | 30 | 1434.0 | 1147 | 2983 | 3948 | 2873 | 10951 | 17 +/-1.8 | 239 | 68 | 249 |
| 10 | 46 | F | 208 | 40 | 11565.0 | 433 | 2941 | 9150 | 2051 | 14575 | 15 +/-4.3 | 229 | 68 | 176 |
| 11 | 30 | M | 208 | 47 | 6989.5 | 3387 | 7843 | 9969 | 6513 | 27711 | 16 +/-3.4 | 141 | 64 | 356 |
| 12 | 47 | F | 208 | 40 | 2468.0 | 1432 | 3145 | 5014 | 4503 | 14093 | 17 +/-4.2 | 188 | 65 | 249 |
| 13 | 57 | F | 26 | 21 | 1455.5 | 736 | 1907 | 2948 | 2176 | 7766 | 12 +/-1.2 | 280 | 68 | 114 |
| 14 | 36 | M | 416 | 60 | 5371.5 | 2682 | 6136 | 9272 | 5869 | 23958 | 24 +/-5.1 | 209 | 62 | 364 |
| 15 | 30 | F | 208 | 72 | 7691.0 | 2296 | 6687 | 9218 | 5867 | 24068 | 21 +/-2.9 | 290 | 67 | 321 |
| Footnotes: a: N, SARS-CoV-2 Nucleocapsid; b: S1, SARS-CoV-2 Spike subunit 1; c: FLS, SARS-CoV-2 Full Length Spike; d: FLT, SARS-CoV-2 Full length Trimer; e: RBD, SARS-CoV-2 Receptor Binding Domain; f: Sum, represents the total of all Spike antigens ; g: iACE2 indicates inhibition of biotinylated ACE2 binding to RBD or FLT antigen complexed to MagPlex beads. add MIA = microsphere immunoassay; MFI = median fluorescence intensity; NAb = NabSure neutralizing antibody; RVP = reporter virus assay |  |  |  |  |  |  |  |  |  |  |  |  |  |  |

We also generated cDBS samples from a commercially available panel of serum samples from a healthy cohort (**Panel E)** as well as a fully vaccinated cohort (**Panel H**). As detailed in the Materials and Methods, serum was combined with an equal volume of human RBCs then spotted (50 μl) onto Whatman 903 filter paper. The spots were allowed to dry at room temperature for 4 h then stored with desiccant at −20°C. Recovery of 90% (±0.04%) total IgG was achieved from a single 3 mm punch (1/8’’) by incubating the punch in 250 μl in TBS with 1% casein for 1 h at room temperature (**Figure 1**). The cDBS eluates were prepared fresh for each assay described below.

**Figure 1.**
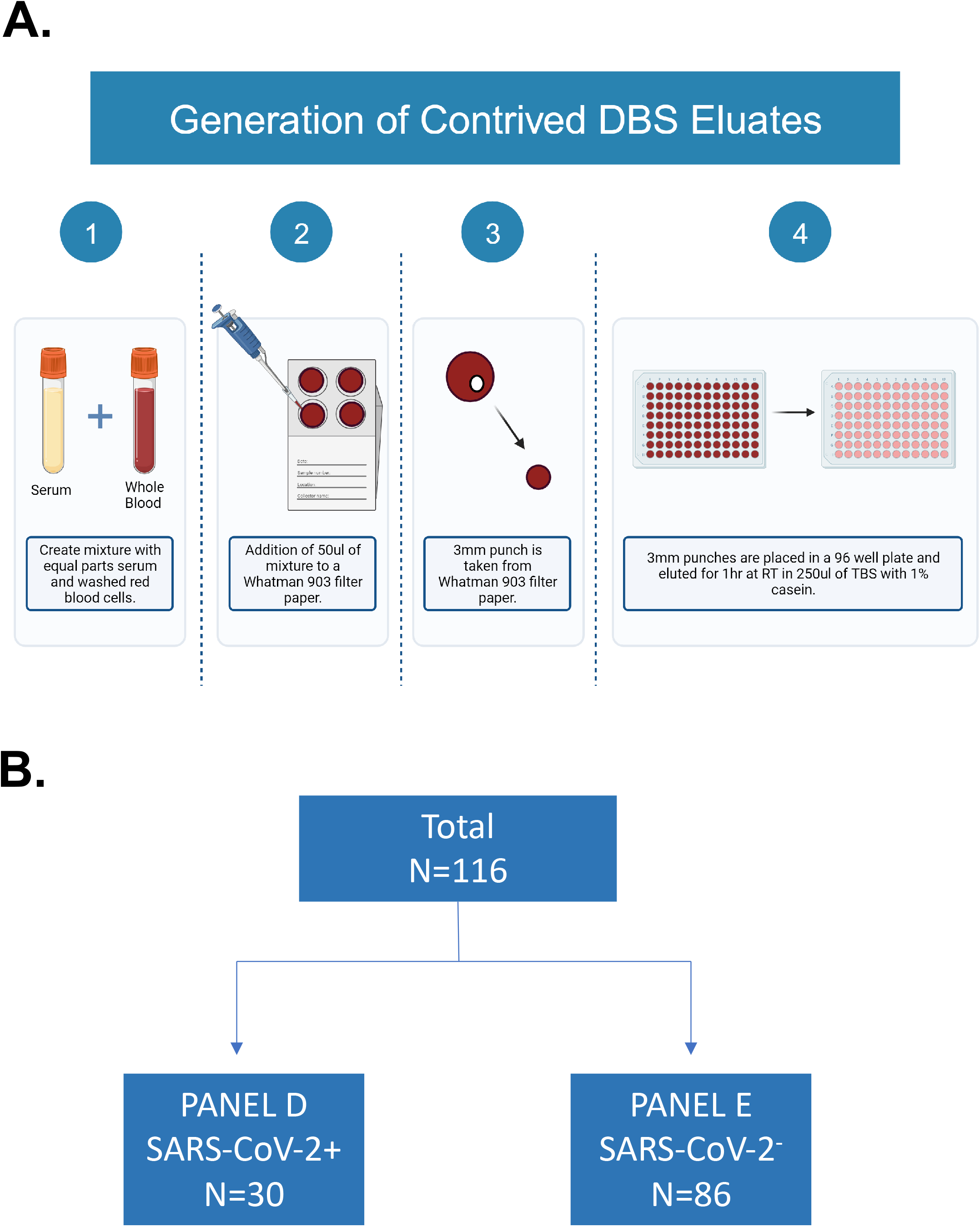
Generation of contrived DBS eluates. (**Top**) Schematic showing generation of contrived DBS (cDBS), DBS punches, and eluate collection. (**Bottom**) Summary of Panels D, E, and H samples used in this study. RT = room temperature, TBS = tris buffered saline

### 3.2 Assessment of SARS-CoV-2 S- and N-specific IgG in DBS eluates by MIA

Using a recently optimized 8-plex MIA targeting different nucleocapsid (N) and Spike antigens, we examined the relationship between MFIs derived from cDBS eluates and PRNT values. Specifically, the 8-plex MIA includes full length Trimer (FLT), full length spike (FLS), S1, RBD and three N antigens as shown in **Table 1**. As expected, Panel D cDBS eluates had higher median fluorescent intensity (MFI) than Panel E cDBS eluates for all seven SARS-CoV-2 antigens (**Figure S1**). In terms of correlation with PRNT (% neutralization @ 1:100 dilution), the values ranged from 0.76 for FLT (p=0.001) and S1 (p=0.001) to 0.82-0.83 for FLS (p =0.0002) and RBD (p=0.0001), respectively (**Figure 2**). Summing the MFIs for all four spike antigens (“sum Spike”) did not greatly increase r values (r = 0.81) (**Figure 2E**). As expected, the correlation between N MFI and PRNT was not significant (r =0.39; p>0.1; **Figure 2F**). We conclude that MFI derived from DBS eluates to a given SARS-CoV-2 spike antigen correlates with virus neutralizing activity as defined by PRNT.

**Figure 2.**
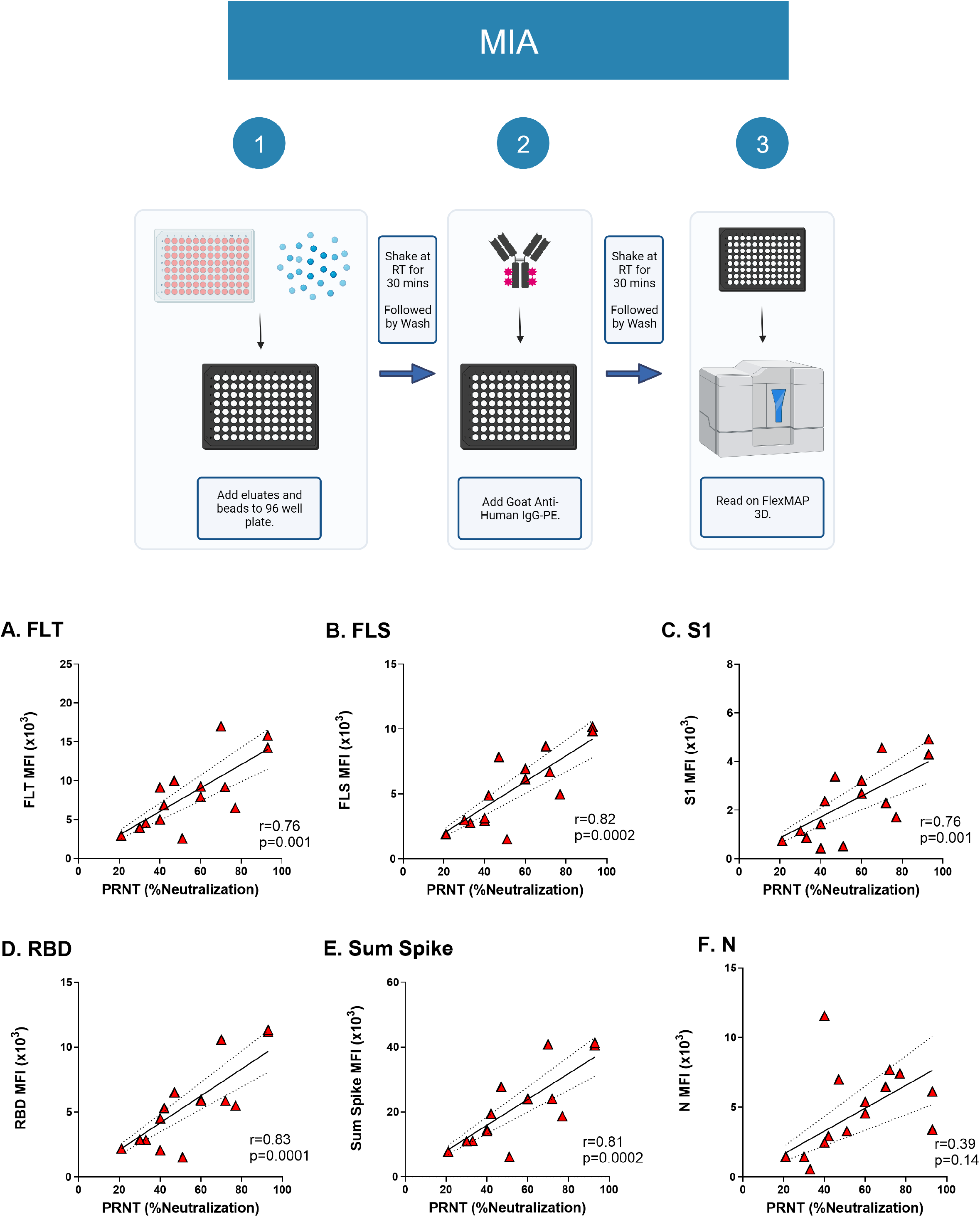
Correlation between PRNT and MIA with cDBS eluates. (**Top**) Workflow for the 8-plex MIA. (**Bottom**) Subset of Panel D cDBS eluates (n=15) underwent testing via 8-plex MIA and were compared to paired serum tested by PRNT. Pearson correlation of MFI values and SARS-CoV-2 neutralizing activity (%) is shown for the following antigens: (A) FLT; (B) FLS; (C) S1; (D) RBD; (E) sum spike (see text for details); and (F) N. The Pearson r value and p-values are shown as insets within each panel.

### 3.3. Pseudovirus neutralization activity in cDBS eluates

Non-replicating, lentiviral-based pseudoviruses expressing SARS-CoV-2 trimeric spike proteins are employed widely to assess SARS and SARS-CoV-2 neutralizing antibody titers in plasma and serum (21–24). However, it is unclear whether DBS eluates are amenable with these cell-based assays. To investigate this, we employed a commercially available non-replicating Reporter Virus Particles (RVP) expressing the SARS-CoV-2 spike protein (D648G) and encoding *Renilla* luciferase (**Figure 3**) (25). We confirmed RVP efficacy using the human monoclonal antibody CC12.3, which neutralizes SARS-CoV-2 with a reported IC_50_ of 0.026 μg/ml (24). In our hands, CC12.3 had an IC_50_ of 0.04 μg/ml (**Figure 3A**). Moreover, the 15 Panel D serum samples had neutralization activities that ranged from 15% to 98% with high concordance with PRNT (r=0.67; p=0.007) (**Figure 3B**). Thus, RVP serves as reliable measure of SARS-CoV-2 activity in the context of serum.

**Figure 3.**
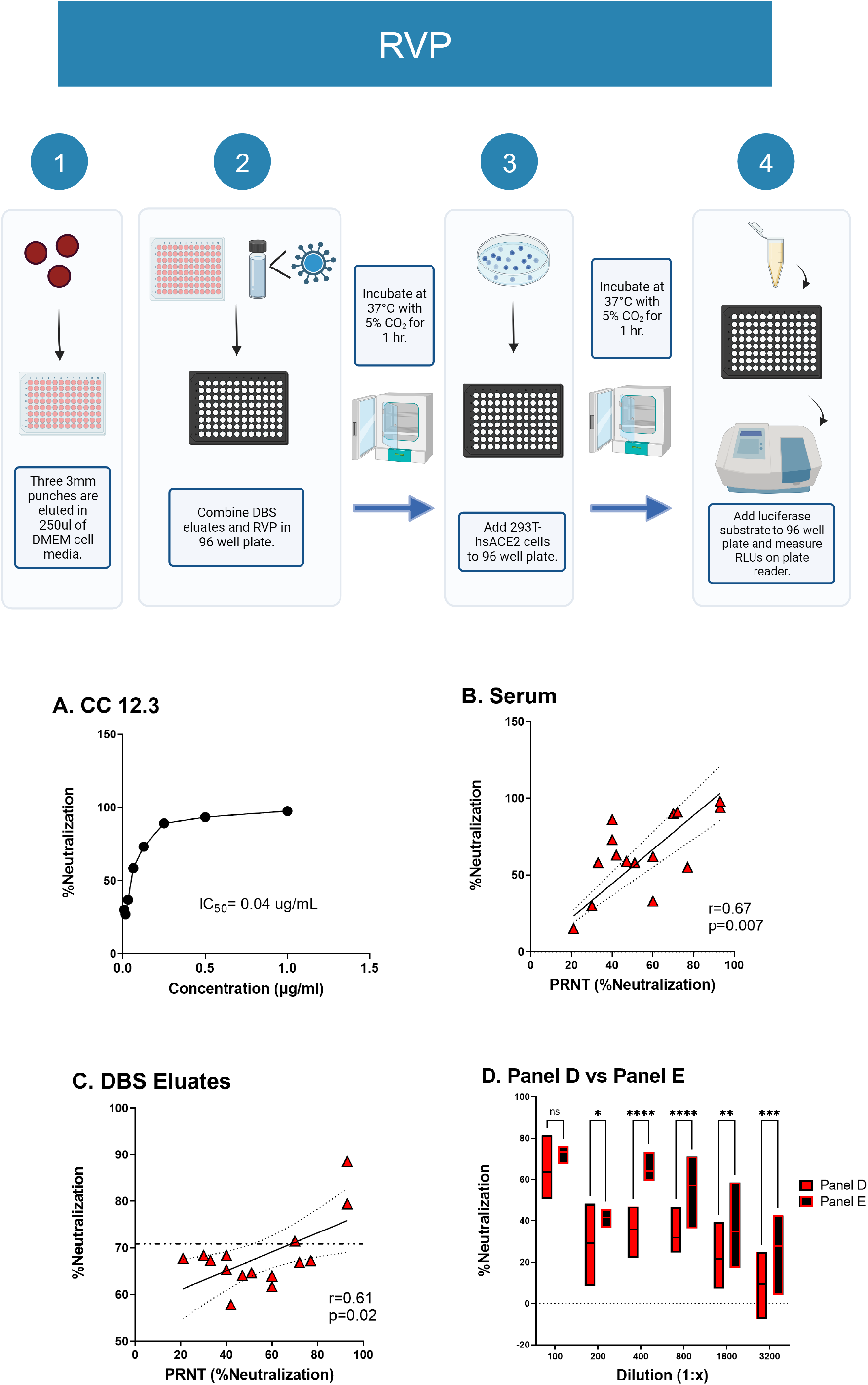
Compatibility of cDBS eluates with RVP neutralizing assay. **(Top)** Workflow for the RVP neutralizing assay. (**Bottom**) (A) Neutralizing titers associated with CC 12.3 with IC_50_ shown as text. (B) Correlation between PRNT and RVP with matched Panel D serum samples; (C) Correlation between PRNT (% neutralization) generated with Panel D serum versus RVP generated from matched cDBS eluates; (D) Comparison of RVP neutralization of Panel D and Panel E cDBS eluates during serial dilutions. Asterisks indicate a significant difference between groups by 2-way ANOVA, where *p=0.02, **p=0.009, ***p=0.0002, ****p<0.0001. For (B) and (C), Pearson r and p-value are shown as a panel inset.

To determine if the RVP assay is compatible with cDBS, we collected eluates from three 1/8” (3 mm) punches from each of the 15 cDBS Panel D spots and 7 Panel E spots as negative controls. Eluates were generated by incubating the punches in DMEM-HEPES at room temperature for 2 h with agitation. Eluates were then diluted 1:50 into DMEM-HEPES medium and then mixed with RVPs prior to application to cells 293T cells constitutively expressing human ACE2 (see **Materials and Methods**).

Panel D eluates had 60-90% neutralization activity [average 68%] that were modestly correlated with PRNT (r value of 0.61; p=0.02) (**Figure 3C**). However, the results were confounded because Panel E eluates (negative controls) had “neutralization” (interfering) activity that was essentially equivalent to Panel D (**Figure 3D**). Thus, RVP assay was not able to distinguish between Panel D and Panel E eluates. Efforts to improve the sensitivity of the assay through dilution (>1:3200), alternative elution buffers, and removal of hemoglobin using reagents designed for this purpose (e.g., HemogloBind™) were unsuccessful (**Figure 3D**; **data not shown**). We conclude that cDBS eluates from convalescent (unvaccinated) individuals have insufficient virus neutralizing titers to distinguish them from unvaccinated controls in the pseudovirus (RVP) platform.

### 3.4 Relationship between PRNT and iACE2

We next tested cDBS eluates for the ability to inhibit ACE2-RBD interactions through use of a Luminex-based assay (iACE2). In the assay, RBD- and FLT-coupled microspheres were incubated with soluble biotinylated-ACE2 in the absence or presence of competitor antibodies (e.g., monoclonal, antisera or DBS eluates), then probed with streptavidin-PE and analyzed by using FlexMap3D (Luminex). A reduction in MFI was interpreted as antibody-mediated inhibition of ACE2 binding to RBD- and FLT. The utility of the in-house iACE2 assay was confirmed using the well characterized monoclonal antibody CC12.3, which neutralizes SARS-CoV-2 by interfering with ACE2-RBD interactions (24). In the iACE2 assay, CC12.3 had an IC_50_ value for WT FLT and RBD of 0.04 ug/mL and 0.02 ug/mL, respectively (**Figure S2**). These values are consistent with those reported by Rogers and colleagues (24).

We examined Panel D DBS eluates in the iACE2 assay, with the expectation that Panel D eluates would have greater inhibition activity than Panel E. Indeed, Panel D had significantly higher iACE2 values (23% average; range 10%-50%) as compared to Panel E (9% average; range 0%-15%) (**Figure 4A**). In terms of the correlation with PRNT, Panel D iACE2 values had an r-value of 0.77 (p<0.001) for RBD-coated beads and 0.72 (p=0.003) for FLT-coated beads. iACE2 (%) values for FLT-coated beads ranged from 25% to 40% (avg 31%) (**Figure 4; Figure S2**). Panel D DBS eluates performed slightly less well than the matched serum samples from which they were derived, at the same dilution, indicating that a degree of ACE2-RBD inhibitory activity is lost during the collection of cDBS eluates. Panel D serum samples (1:100 dilution) had iACE2 activity that correlated with PRNT with r= 0.81 (p<0.001; **Figure S2**). These results demonstrate that iACE2 assay affords a reliable estimate of SARS-CoV-2 neutralization potential using cDBS eluates.

**Figure 4.**
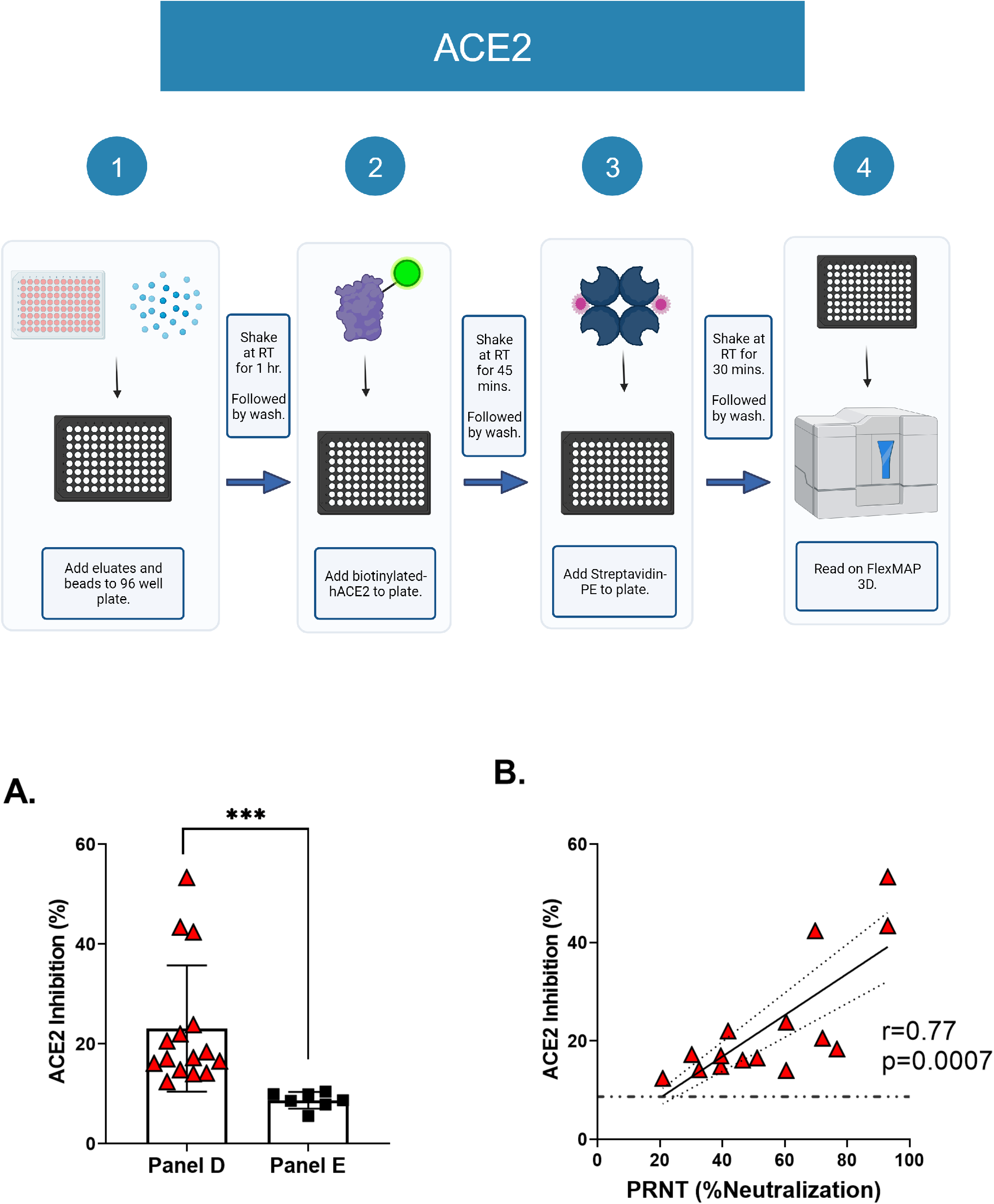
Correlation between Luminex-based ACE2 inhibition and SARS-CoV-2 neutralization activity. (**Top**) Workflow for the ACE2 inhibition (iACE2) assay. (A) Examination of iACE2 activity in Panel D cDBS eluates (n=15) versus Panel E eluates (n=7). In (A), asterisks indicate a significant difference between groups by Welch’s t-test, where ***p=0.0006. (B) Correlation of iACE2 values from Panel D cDBS eluates, as shown in (A), to SARS-CoV-2 neutralizing activity (%) from paired serum analyzed via PRNT. cDBS eluates were evaluated in triplicate. The Pearson r and p-value are shown as a panel inset for (B). The dotted line on (B) represents the average iACE2 activity for SARS-CoV-2 negative cohort (Panel E; n=7).

### 3.5. NAB-Sure^™^

NAB-Sure™ is a commercially available cell-free assay based on successive proximity extension amplification reaction (SPEAR) technology with qPCR as a readout. Disruption of a proximity extension amplification reaction primed by recombinant SARS-CoV-2 S1 protein and human ACE2 by neutralizing antibodies in DBS eluates results in increased CT values. To investigate this technology, we first evaluated the NAB-Sure™ assay using a set of 11 previously authenticated DBS eluates provided by the manufacture and which were subjected to five 3-fold serial dilutions (1 to 243) and tested in duplicate using in-house instrumentation. The NAB-Sure™ assay revealed NT_50_ values that ranged from 210 to 9500 (avg. 2333) with 17.7 %CV (**Figure 5A**). There was a significant correlation between NT_50_ values from DBS eluates tested by NAB-Sure™ and paired serum samples tested by PRNT [Pearson’s r = 0.99; p<0.0001] (**Figure 5A**). Thus, NAB-Sure™ was compatible with DBS eluates and had a high concordance with PRNT.

**Figure 5.**
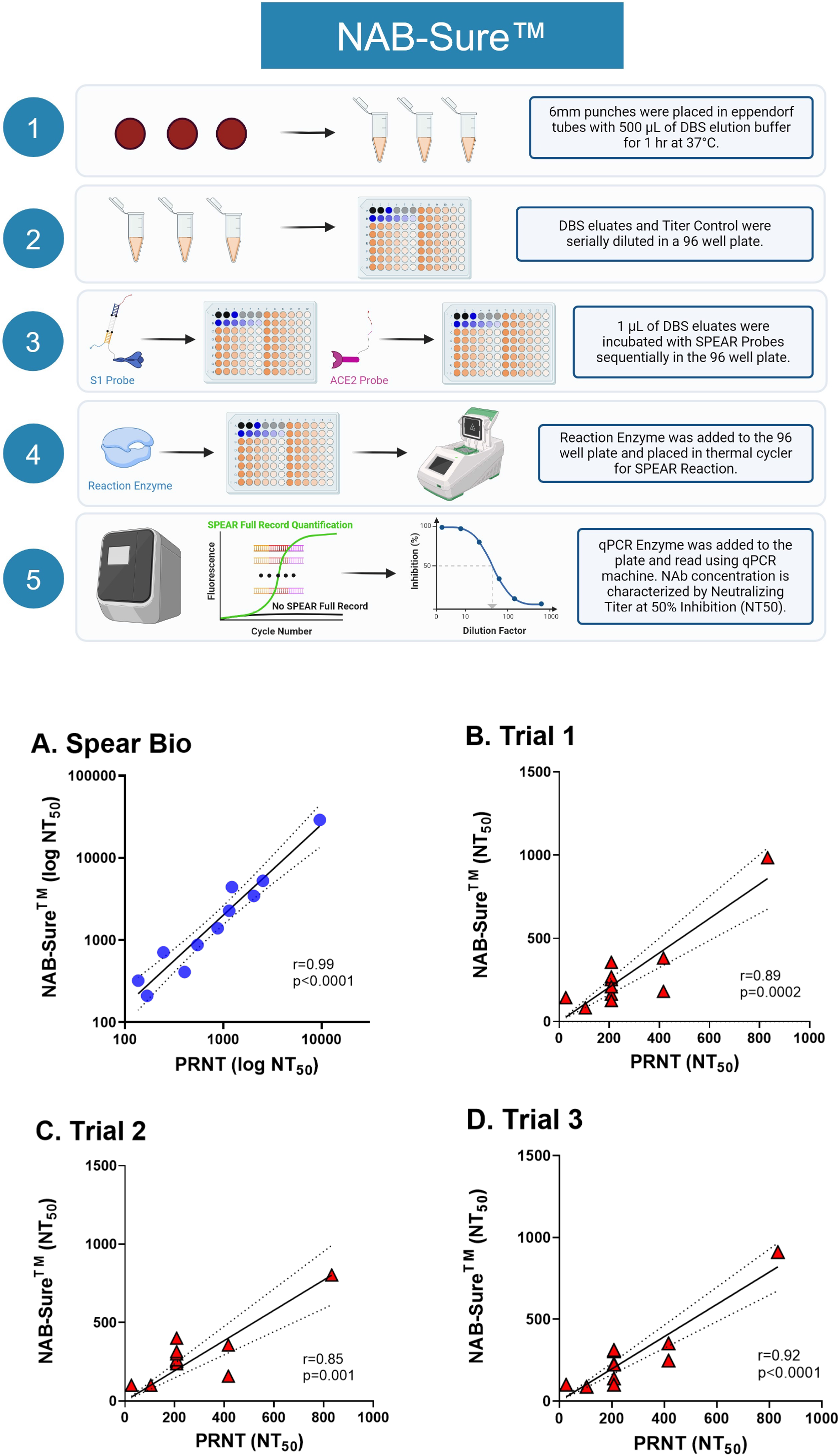
NAB-Sure exhibits high concordance with PRNT. (**Top**) Workflow for the NAB-Sure^TM^ assay. (**Panel A**) Pearson correlation of NAB-Sure ^TM^ NT_50_ from authentic DBS sample eluates compared to PRNT paired serum. DBS samples were evaluated in duplicate. (**Panels B-D**) Pearson correlation comparing NAB-Sure ^TM^ NT_50_ for cDBS eluates and paired serum NT_50_ from PRNT for 14 Panel D samples perfomed in triplicate with each replicate shown. For A-D, Pearson r and p values are in panel insets.

We then used the NAB-Sure™ assay to evaluate cDBS eluates from Panel D (#1-14) and E (#1-7). The samples were tested 3 times in independent reactions under identical assay conditions. Concordance across the different reactions was high, as evidenced by r values between 0.85-0.92 **(Figure 5B-D)**. The average %CV across samples was 20.3%. All reactions reached significance when compared to PRNT. From these results, we conclude that NAB-Sure™ is sufficiently sensitive to detect SARS-CoV-2 neutralizing titers in DBS eluates, based on high degree of concordance with PRNT.

### 3.6 Assessing SARS-CoV-2 neutralizing activity in vaccinated individuals

Studies prior to this point relied on DBS eluates derived from SARS-CoV-2 infected individuals with relatively low overall spike-specific antibody titers as well as SARS-CoV-2 neutralizing levels. With an estimated >90% of the United States population seropositive for SARS-CoV-2 due to vaccination and/or natural infection at this point in time (13), we wished to revisit the utility of DBS eluates in assessing virus-specific antibody titers and neutralizing activity in a vaccinated cohort. For this purpose, we generated cDBS from a commercially available panel (Panel H) of serum samples collected from individuals who had received two doses of the original Moderna (mRNA 1273) vaccine. As shown in **Figure 6**, Panel H DBS eluates (n=15; 15746 ± 1790 MFI) had RBD-specific antibody levels (MFI) what were 1.5 times higher than those in Panel D (10392 ± 3325 MFI) and 44 times higher than Panel E (351 ± 313 MFI). Due to the extremely high MFIs associated with Panel H samples, they were diluted 1:10 for all subsequent assays relative to panels D and E.

**Figure 6.**
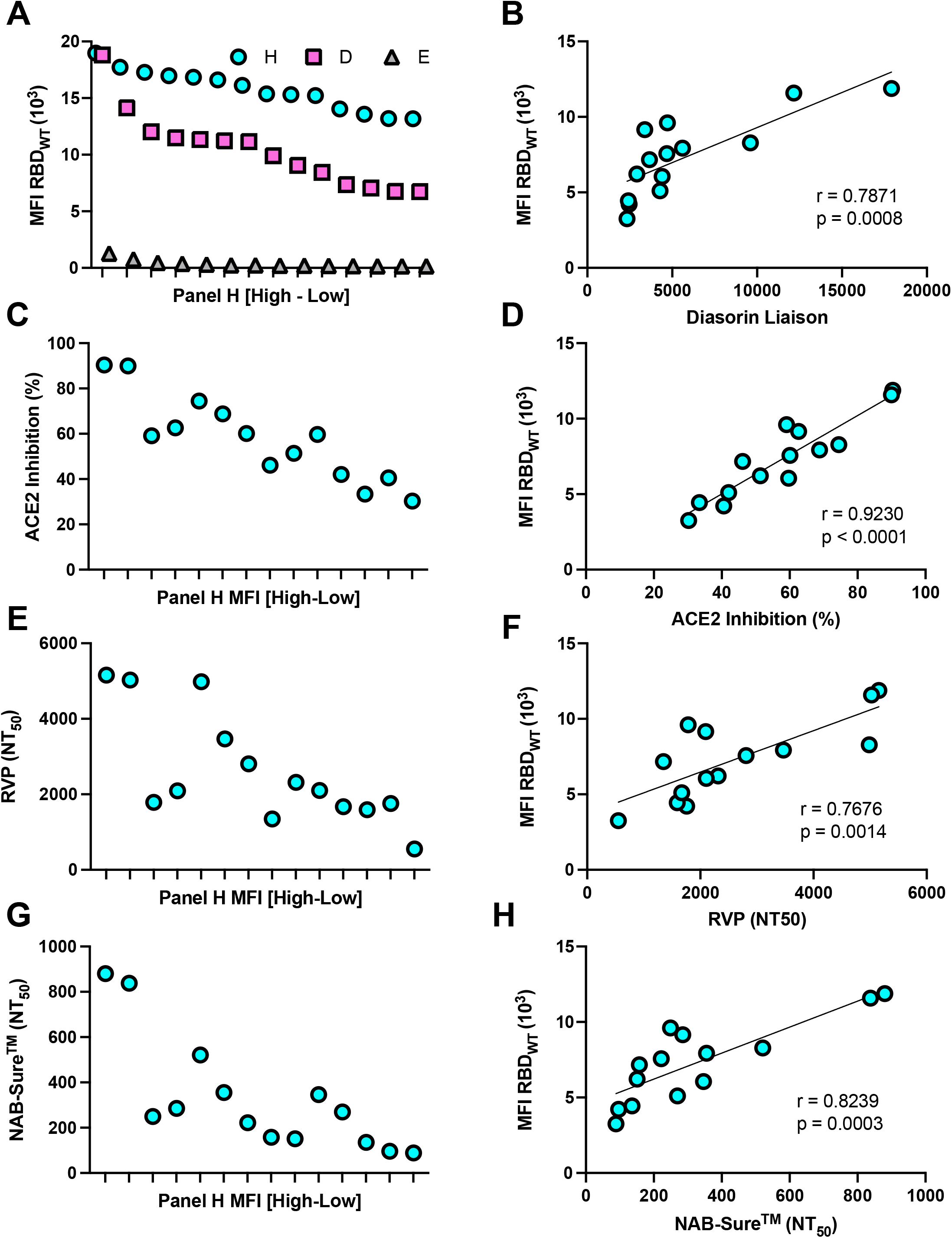
Assessment of SAR-CoV-2 neutralizing titers in cDBS eluates from vaccinated individuals (Panel H). **(A)** Panel H, D and E cDBS eluates were rank ordered from high to low (left to right) based on RBD_WT_ MFI determined in the 8-plex MIA. (**B**) Correlation plot between Panel H RBD_WT_ MFI and Diasorin Liaison values provided from Access Biologics LLC. Panel H cDBS eluate activities in (C) Luminex-based iACE2 assay, (E) RVP neutralization assay and (G) NAB-Sure ^TM^ with corresponding correlation plots shown in right panels (D, F, H). Pearson r and p values are shown as a panel insets. As noted in the text, the Panel H eluates shown in Panel A were diluted 1:10 for the assays descried in panels B-H to avoid saturating MFI values that confound correlation analysis. Moreover, two of the highest Panel H eluates were diluted 1:2 prior to the NAbSure assay to achieve NT_50_ values.

Panel H DBS eluates were evaluated in the iACE2 assay (**Figure 6C, D**), RVP (pseudovirus) neutralization assay (**Figure 6E, F**) and NAB-Sure™ (**Figure 6G, H**). The results demonstrated that all three platforms (iACE2, RVP, NAB-Sure™) were compatible with DBS eluates as evidenced by ACE2 inhibition values approaching 100% (**Figure 6C),** RVP NT_50_ values > 5000 (**Figure 6E**), and NAB-Sure^™^ NT_50_ values approaching 18,000 (**Figure 6G**). There were statistically significant correlations between RBD binding antibody levels (MFI; y-axis) and iACE2 values (x-axis; **Figure 6D),** as well as RBD binding antibody levels (MFI; y-axis) and RVP NT_50_ values (x-axis; **Figure 6F**). This observation is consistent with the earliest observations serology-based correlates of immunity elicited by the spike-encoding mRNA vaccines (14). Moreover, we observed a significant correlation between RBD antibody levels (MFI) and NAB-Sure^™^ NT_50_ values (**Figure 6H**) or RVP NT_50_ values and NAB-Sure^™^ NT_50_ values (r = 0.17; **data not shown**). These results suggest that NAB-Sure™ may perform optimally with DBS eluates samples with lower Spike-specific antibody titers, whereas the iACE2 and RVP assays are better suited for samples with higher Spike-specific antibody titers. Thus, specific assays may be tailored to specific cohorts and their anticipated SARS-CoV-2 titers.

## 4. DISCUSSION

As the World Health Organization (WHO) and other public health agencies integrate long term COVID-19 disease management strategies into the broader landscape of infectious disease, DBS have the potential to serve as a centralized biospecimen type from which to estimate levels of natural and vaccine-induced immunity at population scale (11). Indeed, DBS-based seroprevalence and neutralization assays have been established and deployed to monitor vaccine-induced immunity for the likes of polio (26), human papillomavirus (HPV) (27) and several bacterial pathogens and parasites (28). However, with SARS-COV-2 seroprevalence nearing universality as a result of natural infections/reinfections alongside widespread vaccination, the future utility of DBS to COVID-19 will depend on the development of robust methods to assess functional antibody titers alongside established BAU determinations (14, 29). With this objective in mind, we investigated the compatibility of DBS eluates with different SARS-COV-2 neutralization, pseudovirus neutralization, and ACE2 inhibition assays. While neither the SARS-CoV-2 PRNT nor the pseudovirus (RVP) neutralization assays proved compatible with DBS eluates from convalescent individuals due to high background, values from ACE2-RBD inhibition and NAB-Sure™ assays correlated with neutralization titers (NT_50_ and %) from paired serum. In a cohort of SARS-CoV-2 vaccinated individuals, the ACE2-RBD inhibition and RVP neutralization assays proved more robust than NAB-Sure™, at least under these experimental conditions and with this limited data set. With this in mind, we would recommend tailoring SARS-CoV-2 antibody assessments based on cohorts under investigation. In summary, we conclude that DBS eluates are indeed compatible with multiple SARS-CoV-2 surrogate neutralizing assays and can be applied in parallel with existing and forecasted community and population-based serosurveillance efforts.

Itell and colleagues were the first to investigate the use of DBS to access functional (neutralizing) antibodies from COVID-19 convalescents and SARS-CoV-2 vaccinees (16). In that report, a cell-based SARS-CoV-2 pseudovirus assay proved to be a reliable measure of neutralizing titers with both mock DBS (venous blood spotted on Whatman filter paper) and self-collected fingerprick DBS from vaccinated individuals. In a larger scale study, Roper and colleagues reported success “estimating” SARS-CoV-2 pseudovirus neutralizing titers from fingerprick DBS eluates but noted that multiple samples of the 12 tested were eliminated due to values below limit of detection (30). In their study, fingerprick DBS eluates were not reported at dilutions less than 256, suggesting to us the possibility of interference at low dilutions. They also used eluates from a single 19 mm sample disc into PBS, compared to ∼9 mm (3 x 3 mm) in our study eluted into DMEM and 6 mm by Itell into PBS with 0.1 % Tween-20 (16). Collectively, our results are consistent with DBS eluates being compatible with certain SARS-CoV-2 pseuodovirus assays but only when dilutions exceed ∼256 to eliminate blood-based interference.

McDade and colleagues reported the compatibility of DBS eluates with an ACE2-RBD inhibition (iACE2) assay on the Mesoscale Diagnostics platform (17). We now extend those results and demonstrate that both an in-house Luminex-based assay and a commercial real-time PCR-based assay (NAB-Sure™), give rise to inhibition values that strongly correlate with neutralization titers derived from PRNT. While we recognize that the iACE2 assay is strictly a surrogate neutralization assay, both platforms described herein have the potential to be integrated alongside existing high-throughput SARS-CoV-2 multiplexed serologic assays. At present, we employ an 8-plex SARS-CoV-2 microsphere immunoassay (MIA) run in 384 well format on the Luminex platform with automated liquid handling (12). The in-house iACE2 assay described here is also run on the Luminex platform using the same magnetic microspheres as the MIA. The overall costs of the iACE2 assay are at least an order or magnitude less expensive than cell-based pseudovirus assay with commercial RVPs. Moreover, the in-house iACE2 assay is easily multiplexed to include microspheres coated with RBD from circulating VoCs and Omicron subvariants side-by-side with ancestral RBD. Indeed, we have applied a 4-plex RBD assay to several thousand primary DBS samples collected from an elderly cohort (G. Mirabelle, K. Berman, N. Mantis, unpublished results).

The NAB-Sure™ assay has the advantage of being ultrasensitive and amenable to scale-up and automation and provides NT_50_ titers similar to PRNT. Moreover, as NAB-Sure™ is a commercial kit, there were no in-house costs associated with reagent quality assurance or validation making the kit comparable (cost wise) with most pseudovirus assays. In our hands, NAB-Sure™ proved most robust with DBS eluates in which SARS-CoV-2 levels were on the low end (e.g., convalescent samples), while RVP and iACE2 performed well with high titer samples associated with vaccination. It is unclear whether this distinction will apply to DBS samples aimed at evaluating antibody titers against SARS-CoV-2 variants of concern (VoC) like the highly divergent JN.1. In fact, a tiered approach to interrogating DBS eluates may be in order where MFI values alone provide a rough idea of neutralizing titers within a given sample especially when dealing with thousands of samples from diverse cohorts (31). In terms of functional antibodies, the Luminex-based iACE2 assay is both affordable and adaptable to any given RBD. In fact, we have performed iACE2 multiplexing with four different RBD variants simultaneously on hundreds of primary DBS samples from multiple different cohorts (G. Mirabile,K. Berman, N. Mantis, *manuscript in preparation*). Ultimately, both the NAB-Sure™ and pseudovirus assays can be employed as a means to establish quantitative NT_50_ equivalents not achievable with MFI or iACE2 alone.

The use of DBS for infectious disease serology and functional antibody testing is not without intrinsic challenges, including variability in blood collection by fingerpick, uniformity of antibody recovery in eluates, and limited sample volumes (11). Nonetheless, the benefits of DBS as a biospecimen type for surveillance of hard-to-reach communities are well recognized. For example, in work spearheaded by the CDC, DBS from ∼1,000 children in Bangladesh were sampled for neutralizing antibodies to polio types 1, 2, and 3 using a modified poliovirus microneutralization assay (26). In the case of polio, a neutralizing antibody titer of 1:8 correlated with protection and was used as the threshold definition of seropositivity. Similarly, in a study examining natural versus vaccine-induced antibody responses to human papillomavirus (HPV), neutralizing activity was detected in 2 mm DBS punches eluted into 0.2 mL saline (27). Another consideration is that for SARS-CoV-2 at least there is a strong congruence between oral fluids and DBS (K. DeRosa, N. Pisanic, K. Kruczynski, L. Styer, M. Parker, C. Heaney, and N. Mantis, *manuscript in preparation*). Thus, DBS and oral fluids might potentially be used interchangeably as means to survey large cohorts for immunity and vaccine responsiveness (32).

## ACKNOWLEDGEMENTS

We gratefully acknowledge SpearBio, Inc (Woburn, MA) for providing matched serum and DBS samples as well as assistance in optimizing NAB-Sure™ assay at the Wadsworth Center. The SARS-CoV-2 reagents provided by MassBiologics (Boston, MA) were generated under a Project Award Agreement to NJM from the National Institute for Innovation in Manufacturing Biopharmaceuticals (NIIMBL), United States and financial assistance award 70NANB20H037 from the US Department of Commerce, National Institute of Standards and Technology. Research reported in this publication was supported by the National Cancer Institute of the National Institutes of Health under Award Number U01CA260508. The content is solely the responsibility of the authors and does not necessarily represent the official views of the National Institutes of Health.

## CRediT

| Author | Conceptualization | Curation | Formal analysis | Funding acquisition | Investigation | Methodology | Admin | Resources | Software | Supervision | Validation | Visualization | Writing-original | Writing-review |
| --- | --- | --- | --- | --- | --- | --- | --- | --- | --- | --- | --- | --- | --- | --- |
| KB |  |  | x |  | x | x |  |  |  |  | x | x | x | x |
| GVS |  |  |  |  | x | x |  |  |  |  |  |  |  |  |
| HN |  |  |  |  | X | X |  |  |  |  |  |  |  |  |
| JMR |  |  | X |  | X | X |  |  |  |  | X | X | X | X |
| RB |  |  | X |  | X | X |  |  |  |  | X | X | X | X |
| AKD |  | X | X |  |  |  |  |  |  |  | X |  | X | X |
| KLD |  | X |  |  | X | X |  |  |  |  |  |  |  | X |
| GM |  |  |  |  | X | X |  |  |  |  |  |  |  |  |
| RCG |  |  | X |  | X | X |  |  |  |  | X |  |  |  |
| APD |  |  |  |  | X | X |  |  |  |  | X |  |  |  |
| KAM |  |  |  |  |  |  | X | X |  | X |  |  |  | X |
| MMP | X |  | X |  |  |  | X | X |  | X |  |  |  |  |
| LMS | X |  | X |  |  |  | X | X |  | X |  |  | X | X |
| NJM | X |  | X | X |  |  | X | X |  | X |  |  | X | X |

## Supplemental Figure Legends

**Figure S1.**
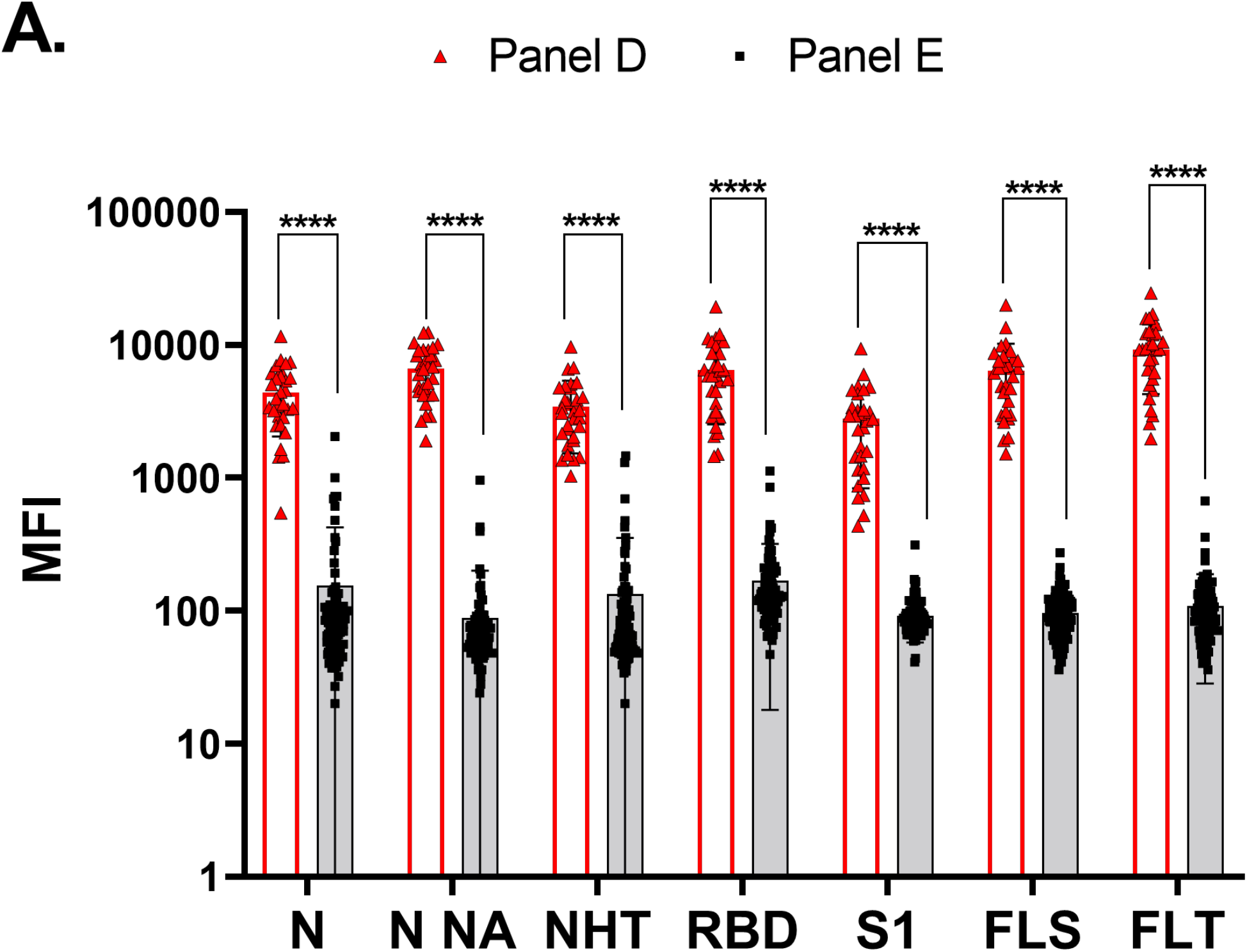
Comparison of Panel D and Panel E cDBS. The full cohort for Panel D (n=30) and Panel E (n=86) samples underwent testing via 8-plex MIA. MFI values for the following antigens are compared: N, N NA, NHT, RBD, S1, FLS and FLT. Asterisks indicate a significant difference between groups by 2-way ANOVA, where ****p<0.0001.

**Figure S2.**
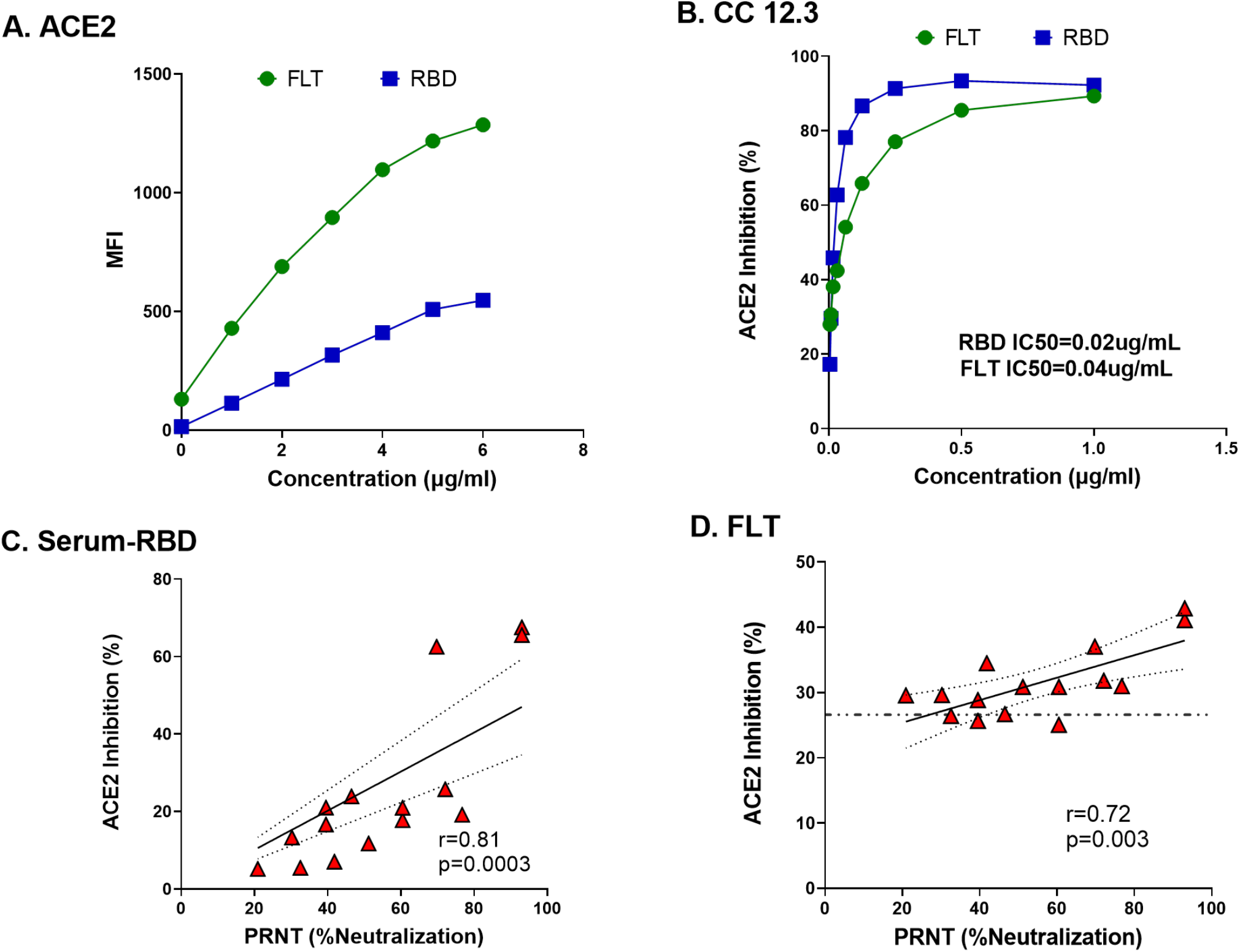
Validation of Luminex-based ACE2 inhibition assay. (A) MFI values (binding) was assessed in the context of either FLT or RBD with varying concentrations of biotinylated hACE2. (B) ACE2 inhibition titers associated with CC 12.3 in the context of FLT or RBD. IC_50_ is shown as text. (C) Pearson correlation of Panel D serum analyzed by ACE2 inhibition and SARS-CoV-2 neutralizing activity (%), r-value and p-value shown as text. (D) Pearson correlation of Panel D DBS eluates analyzed by ACE2 inhibition and SARS-CoV-2 neutralizing activity (%), in the context of FLT. The r-value and p-value are shown as text. The dotted line on (D) represents the average iACE2 activity for SARS-CoV-2 negative cohort (Panel E; n=7).

